# Spatial and temporal control of CRISPR/Cas9-mediated gene editing delivered via a light-triggered liposome system

**DOI:** 10.1101/725465

**Authors:** Yagiz Alp Aksoy, Wenjie Chen, Ewa M Goldys, Wei Deng

## Abstract

The CRISPR-Cas9 and related systems offer a unique genome editing tool allowing facile and efficient introduction of heritable and locus-specific sequence modifications in the genome. Despite its molecular precision, temporal and spatial control of gene editing with CRISPR-Cas9 system is very limited. We developed a light-sensitive liposome delivery system that offers a high degree of spatial and temporal control of gene editing with CRISPR/Cas9 system. We demonstrated its high transfection efficiency, by assessing the targeted knockout of eGFP gene in human HEK293 cells (52.8% knockout). We further validated our results at a single-cell resolution using an in vivo eGFP reporter system in zebrafish (77% knockout). To the best of our knowledge we reported the first proof-of-concept of spatio-temporal control of CRISPR/Cas9 by using light-triggered liposomes in both *in vitro* and *in vivo* environment.

## INTRODUCTION

Control of gene activity is one of the principal methods to study details of physiological processes at a cell level in live organisms, and therefore it holds the key to the understanding of the machinery of life. The CRISPR (Clustered Regularly Interspaced Short Palindromic Repeats) and related methods revolutionised this field by introducing an efficient platform for precise engineering of genes ^1, 2^. In this approach, a nuclease protein (Cas9) introduces a double-stranded break in the target sequence of a DNA molecule, enabling the incorporation of a new sequence into the genome as directed by the guide RNA (gRNA) repair template. A truly transformative technology, CRISPR makes it possible to explore gene functions by facile knocking out genes, adding transgenes, or by programmable transcriptional and post-transcriptional regulation ^3-6^. Major advances have recently been made in the clinical applications of CRISPR ^7, 8^ through the development of therapeutics that can specifically disrupt the expression of disease-relevant genes ^8-12^. So far, CRISPR has been successful in cancer CAR-T immunotherapy and to treat primary defects of the immune system, hemoglobinopathies, hemophilia, metabolic disorders, and muscular dystrophy ^6, 10, 13^. CRISPR/Cas9 gene editing has been used to reprogram cells including iPS (Induced Pluripotent Stem) cells ^14^, and to prevent/treat viral and bacterial infection, by targeting genes conferring virulence or antibiotic resistance ^15^. The applications of this system have also been extended to other fields, including biotechnology and agriculture ^16, 17^.

The safe and efficient delivery of CRISPR–Cas9 components to targeted cells and tissues remains one of the key challenges for successful gene editing ^9^. The most widely reported approach for proof-of-principle studies is transfection of plasmid DNA carrying nuclease and gRNA expression cassettes. This method is unsuitable for clinical translation due to low transfection rates, DNA-related cytotoxicity, and the possibility of random integration of plasmid fragments including bacterial DNA sequences into the genome ^10^. Viral gene transfer is the current leading approach for CRISPR delivery in clinical trials, with lipid nanoparticles and physical methods such as electroporation also clinically relevant ^6^. In viral delivery, both components of CRISPR, the guide RNA and Cas9 are introduced into cells and tissues by lentiviral, adeno-viral, or adeno-associated viral vectors (AAV) created using recombinant DNA technology^7, 8, 10, 15^. The viral expression vectors offer a limited level of spatial and temporal control over the gene editing process, and the possibility of unintended biological consequences. This is because once the nuclease gene is delivered, there is no intrinsic mechanism to terminate the replication. As a result nuclease expression may be long-lasting or excessive leading to suboptimal efficiency of genome editing, or adverse immune response ^18^. Moreover, some therapeutic genes are too large to be readily packaged and transferred by available delivery viral vectors (4.7 kB for AAVs) and need special approaches to achieve transient nuclease expression ^10^. Physical methods are applicable to *ex vivo* gene editing, (e.g. electroporation of mRNA encoding the nucleases and gRNAs is a preferred method to edit T cells and hematopoietic stem cells), however they are unsuitable *in vivo* ^10^.

Among various viral-free nanoformulations, lipid, polymer and other nanomaterials have been explored to deliver the CRISPR-Cas9 systems for therapeutic purposes or to establish knockout animal models ^19, 20^. For example, Cas9 mRNA and the sgRNA have been loaded onto lipid nanoparticles and delivered to murine liver with high efficiency ^20-22^. Nanoparticle-mediated delivery of a single CRISPR component spCas9 mRNA has been used in combination with AAVs encoding a sgRNA and a repair template with the efficiency of over 6% ^23^. Furthermore, modified nanoparticles have been loaded with a donor template to achieve homology-directed repair ^24^. The nanoparticle platforms have also shown to accommodate multiple components of CRISPR–Cas9 into a single carrier ^20^.

Liposomal formulations represent an alternative option for nanomaterial-based gene delivery, well established in earlier literature including by our group ^25-30^. Liposomes have high loading capacity, can carry complex cargos and their biodistribution and pharmacokinetics can be refined through sophisticated multifunctional formulations. They also offer chemically defined compositions and high level of control over drug delivery, including externally triggered or inducible, self-regulating control over drug release ^26, 31-33^. Our earlier work on triggered liposome delivery systems demonstrated their capability in carrying and releasing short DNA ^30^, plasmid DNA ^28^ and chemotherapy drug ^26^ *in vitro* and *in vivo*. Liposomes carry no risk of genomic integration and unintended immune activation can be easily avoided. The dose control available in nanoparticle delivery is also important for CRISPR/Cas9 gene editing, as the duration and magnitude of nuclease expression has been found to be a critical parameter for the level of both on-target and off-target nuclease activity ^10^. Liposomal carriers were previously utilized to deliver CRISPR/Cas9 system to cultured cell lines and animal models ^27, 34-38^. However, like lipid-based nanocarriers, simple liposomes only able to passively release CRISPR/Cas9, without spatial or temporal control over gene editing.

New generation liposomes offer the option of on-demand payload release by external or internal stimuli such as light pH, temperature etc ^39-41^. External light source is a convenient stimulus employed in activation of on-demand release from the liposomes due to easily adjustable spectral properties, illumination intensities and times. Furthermore, spatial and temporal control of light sources provides an extra benefit to precisely tune the cargo release.

In this work we realised light-triggered liposomes by incorporating a photosensitive molecule, verteporfin (VP) inside the lipid bilayer. Under light illumination at 690 nm wavelength, VP reacts with available oxygen molecules and generates singlet oxygen which oxidises the unsaturated lipid components and leads to destabilisation of the liposomal structure and CRISPR release (Fig.1A). We demonstrated temporal and spatial control of CRISPR gene *in vitro* and *in vivo.* Efficient GFP gene transfection was carried out using light-triggered liposomes encapsulating Cas9-gRNA Ribonucleoprotein (RNP) in human kidney cells (HEK293) and in a zebrafish model. The zebrafish is a well-established system to model human disease ^42^, and many cellular pathways are highly evolutionarily conserved between humans and zebrafish. In addition, zebrafish have a fully sequenced genome in which 82% of human disease genes have clear homologs ^43^. The vast genetic toolbox available for the manipulation of zebrafish allow forward- and reverse-genetic screen studies while its transparent embryos make zebrafish an excellent organism for *in vivo* imaging ^42^.

**Figure 1.**
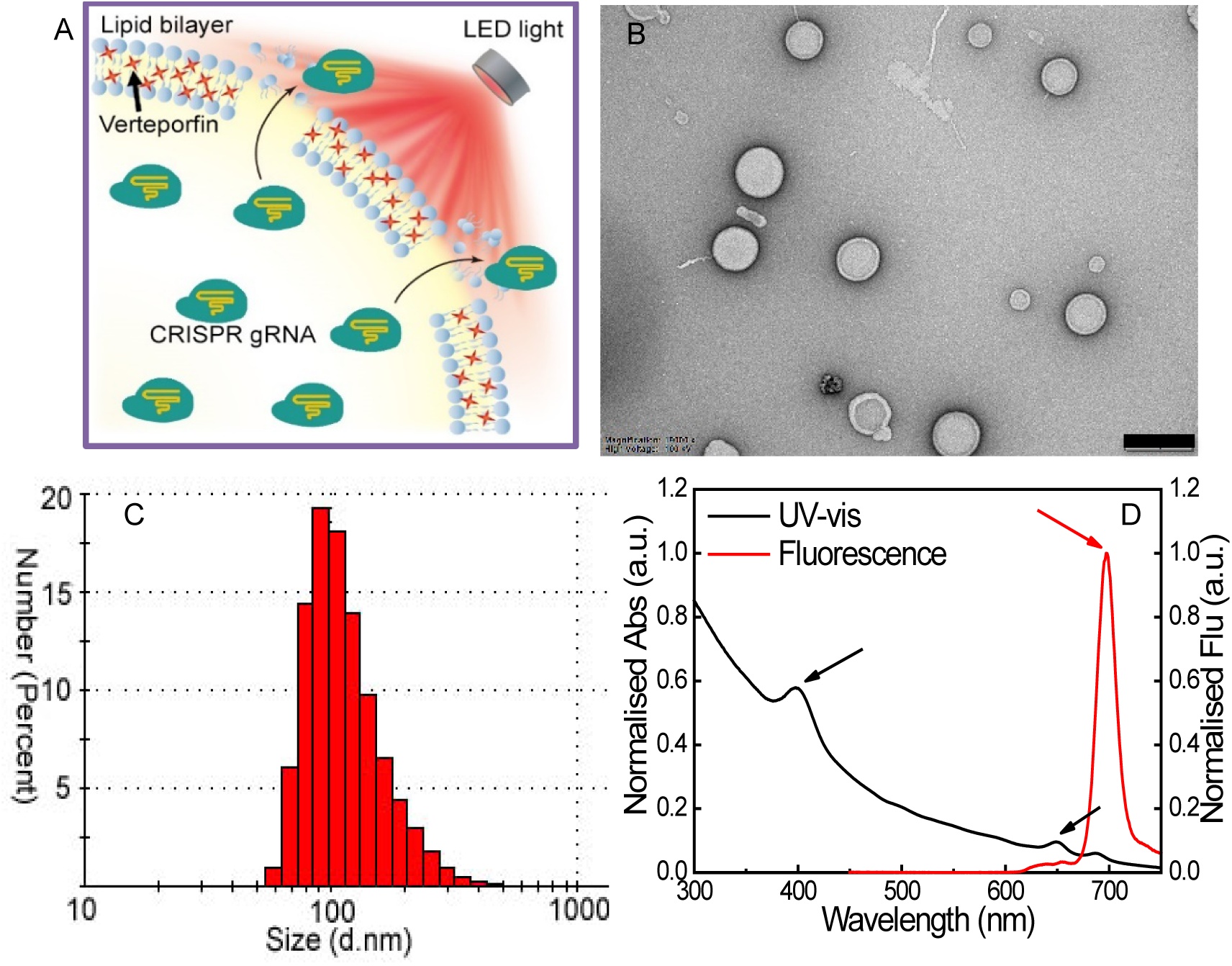
(A) Schematic illustration of the release of CRISPR agents from a light-triggered liposome under a LED light source at 690 nm; (B) The typical TEM image of liposomes incorporating verteporfin (scale bar = 500 nm); (C) Size distribution of liposome suspension; (D) Absorption and fluorescence spectra of verteporfin loaded inside liposomes. Black and red arrows indicate the characterized peaks of verteporfin.

We complexed the liposomes with Cas9 RNP and microinjected the mixture solution into zebrafish embryos expressing eGFP gene, followed by light illumination at 690 nm. To establish a simple and quantitative readout for gene knockout we focused on the large slow-muscle cells in the zebrafish trunk. Zebrafish slow-muscle is a single layer of parallel fibers that encase the fish beneath the skin, rendering them accessible to rapid and accurate quantitation by fluorescence microscopy. In this work we used a double transgenic zebrafish strain that expressed eGFP under the control of the slow-muscle smyhc1 promoter. To evaluate the efficiency of the sgRNAs, we targeted a region in eGFP and confirmed the loss of eGFP fluorescence in individual slow-muscle cells at 72 hours post-fertilization (hpf)

## MATERIALS AND METHODS

Lipids (DOTAP and DOPE) were purchased from Avanti Polar Lipids (Alabaster, AL, USA). Verteporfin, cholesterol (Chol) and chloroform were purchased from Merck Australia. Dulbecco’s modified Eagle’s medium, fetal bovine serum, trypsin, optiMEM, Dulbecco’s Phosphate-buffered saline, Truecut cas9 v2, GFP gRNA and lipofectamine were purchased from ThermoFisher Australia. Zyppy Plasmid MiniPrep Kit was purchased from Zymo Research. MEGAshortscript T7 kit and mirVana miRNA isolation kit were purchased from Invitrogen Australia. Cas9 protein used *in vivo* experiments was obtained from Toolgen, Inc.

### Preparation of liposomes incorporating Cas9-gRNA RPN

The liposome formulation was prepared based on our previous method with minor modification ^26^. Briefly, DOTAP, DOPE, Chol and verteporfin at mole ratio of 1:0.94:1:0.06 were mixed 500 µL chloroform. The mixture solvent was then evaporated under argon gas stream. The thin lipid film was formed around the wall of the test tube and hydrated with DI water by vigorous stirring for 30 min until the suspension was homogenized. For preparation of liposomes incorporating Cas9 gRNA RPN, the lipid film was fully resuspended in 500 µL DI water solution containing sgRNA (1 mg mL^-1^) and Cas9 protein (5 mg mL^-1^). The hydrated liposome suspension was extruded 11 times through a 200 nm polycarbonate membrane in a mini-extruder. The absorption and fluorescence spectra of the liposomes incorporating VP were measured with a UV-VIS spectrometer (Cary 5000, Varian Inc.) and a Fluorolog-Tau3 System (HORIBA Scientific) with 425 nm Xe lamp excitation, respectively.

### Characterization

The zeta potential and size distribution of liposome samples were determined by DLS using a Zetasizer 3000HSA. After 2 min balance at 25°C, each sample was measured in triplicate and data were collected as the mean ± standard deviation (SD). Prior to transmission electron microscopy (TEM) imaging of liposome sample, the TEM grid specimens were prepared using the negative staining method. Briefly, a copper grid was placed onto a drop of 10 µL liposome suspension, allowing the grid to absorb samples for 3 min, followed by staining with 2% (w/v) phosphotungstic acid for another 3 min. After air-drying of the sample overnight, the grid specimens were then imaged using a TEM (Philips CM 10) with an acceleration voltage of 100 KV. Images were captured with the Olympus Megaview G10 camera and processed with iTEM software. The absorption and fluorescence spectra of liposomes and pure VP were measured with a UV-VIS spectrometer (Cary 5000, Varian Inc.) and a Fluorolog-Tau3 System (HORIBA Scientific) with 425 nm Xe lamp excitation, respectively.

### Assessment of *in vitro* GFP gene transfection via light-triggered liposomes

A transgenic HEK293 containing GFP gene in the genome (ThermoFisher via MTA) was used in cell experiments. They were grown in DMEM containing 10% fetal bovine serum and 1% antibiotics. Before transfection, HEK293 were seeded on a 24-well plate at the density of 1×10^5^ cells/well, followed by overnight incubation. 1 ml of optiMEM solution containing 100 µL liposomes incorporating Cas9 gRNA RPN was added to each well. After 2 hr incubation, the old medium was replaced by the fresh one, followed by illumination of LED light (0.15 mW/cm^2^) at 690 nm for 2 min, 4 min and 6 min, respectively. After the treatments, the cells were incubated for another 48 h. The GFP fluorescence signal from the cells was imaged using under a FV3000 confocal laser scanning microscope. A laser at 488 nm was used for GFP excitation. Quantitative analysis of GFP signal was conducted by using ImageJ software, which indicated GFP gene knockout efficacy under different experimental conditions.

### Zebrafish Embryos

Zebrafish embryos and adults were maintained and handled according to zebrafish facility SOPs, approved by Animal Research Ethics Committee at Macquarie University and in compliance with the Animal Research Act, 1985 and the Animal Research Regulation, 2010. Adult zebrafish were maintained under standard conditions. *smyhc1:eGFP* line was generated from *Acta1:eBFP2;smyhc1:eGFP* line ^44-46^.

### Target Site Selection

For the initial screen, CRISPR sgRNA target sites were selected manually within the early 5’ region of eGFP gene that match the sequence GN18GNGG according to Ref. ^47^. To avoid any off-target effects, these sites were checked for uniqueness in BLASTN (Zv9) using Bowtie and Bowtie2 methods, and the pre-defined specificity rules that do not tolerate any mismatch in the first ten 3’ bases of the target site.

### Production of sgRNA

To generate templates for sgRNA transcription, target gene-specific complementary oligonucleotides containing the 20 base target site without the PAM, were annealed to each other, then cloned into a plasmid containing T7 promotersequence and tracrRNA tail. The resulting sgRNA template was purified using Zyppy Plasmid MiniPrep Kit. For making CRISPR sgRNA, the template DNA (from the step above) was first linearized by BamHI digestion, then purified using a QIAprep column. CRISPR sgRNA is generated by *in vitro* transcription using MEGAshortscript T7 kit. After *in vitro* transcription, the sgRNA (∼140 nucleotides long) was purified using mirVana miRNA isolation kit. The size and quality of resulting sgRNA was confirmed by electrophoresis through a 3%(wt/vol) low-range agarose gel.

### Microinjection of liposome-Cas9 RNPs into zebrafish embryos

On the day of injections, the injection mix was prepared as indicated in Table 1.

**Table 1.**
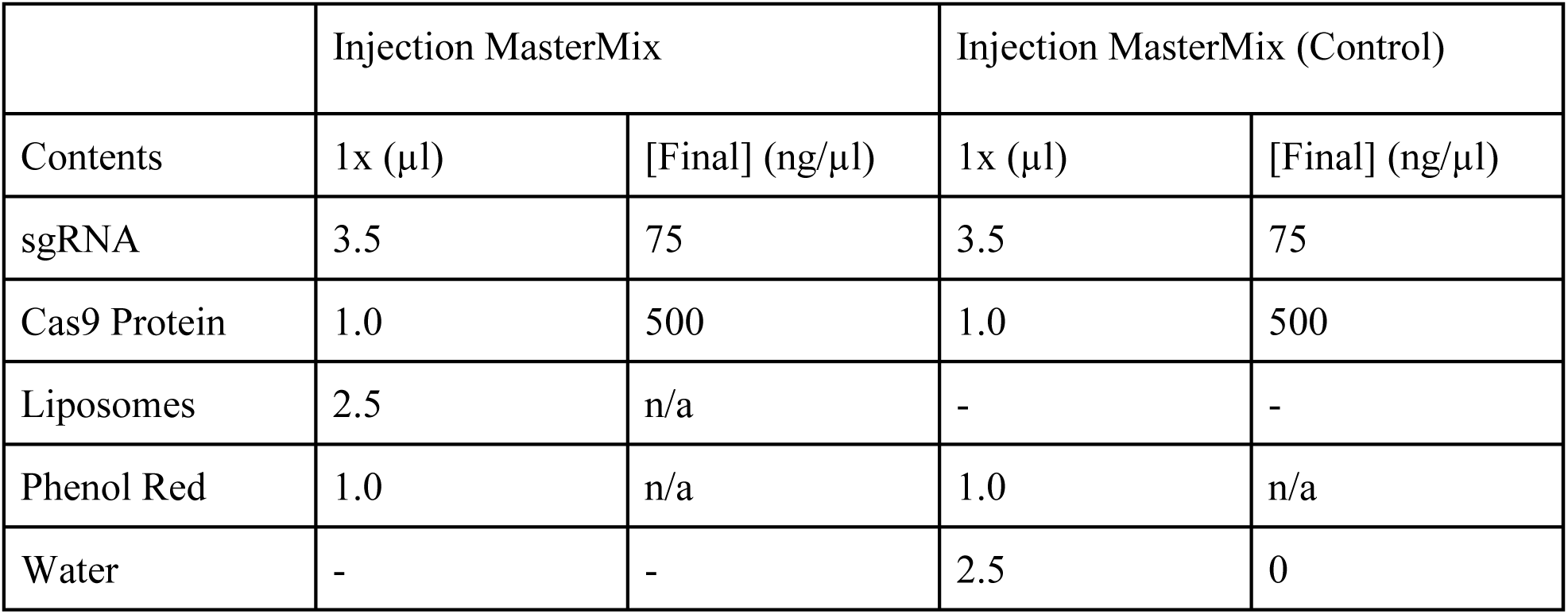
Volume of each agent used for microinjection

|  | Injection MasterMix |  | Injection MasterMix (Control) |  |
| --- | --- | --- | --- | --- |
| Contents | 1x (μl) | [Final] (ng/μl) | 1x (μl) | [Final] (ng/μl) |
| sgRNA | 3.5 | 75 | 3.5 | 75 |
| Cas9 Protein | 1.0 | 500 | 1.0 | 500 |
| Liposomes | 2.5 | n/a | - | - |
| Phenol Red | 1.0 | n/a | 1.0 | n/a |
| Water | - | - | 2.5 | 0 |

For the initial screen, zebrafish TAB WT embryos were collected. Injection components were mixed and incubated at room temperature for 5 min to form complex, then stored on ice. The injection mastermix was loaded into the needle and microinjected into zygotes using standard zebrafish injection protocols. Delivery of 2 nl of injection mixture into the single cell (not the yolk) aimed. The injected eggs were grown in 1x egg water in 100mm plastic petri dish and kept in the incubator at 28°C. Embryo density did not exceed more than 60 embryos in 25 mL egg water per petri dish. Some uninjected embryos (control group) were kept from the same clutch and grown at 28 °C. Embryos were grown to 48-72hpf.

### *In vivo* recombination analysis

Embryos with developmental defects were sorted out at the end of 24 hpf, 48 hpf and 72 hpf. Only morphologically normal looking embryos were kept. Approximately 70-80% of embryos appear normal at 72 hpf. At 72 hpf, 16 embryos were randomly selected and anesthetised using Tricane. Anesthetised fish were mounted on 1% low-melting agarose in glass bottomed 35mm Petri dishes. The trunk of mounted embryos was screened for eGFP signal using Leica DMi3000 inverted microscope.

### Microscopy

Images were taken on a Leica DMi3000 inverted microscope and Zeiss confocal microscope. Fish embryos were embedded on 1% low-melting agarose in 35mm glass bottom petri dish. Sections were focus-stacked using Zerene Stacker software. Virtual cross sections of the fish embryos were generated and analysed using Imaris software.

## RESULTS

### Characterization of liposomes

Fig.1B shows the typical TEM image of liposomes loaded with verteporfin. The size distribution and zeta potential of liposomes were confirmed by dynamic light scattering, with an average size of about 167.5 +/-1.9 nm and surface charge of to 28 ± 1.1 mV (Fig.1C). The absorption and fluorescence spectra of verteporfin loaded inside liposomes were demonstrated in Fig. 1D, where the characterised peaks of verteporfin were clearly observed, as indicated in the figure.

### Assessment of *in vitro* GFP gene knockout by using liposome-CRISPR/Cas9

The confocal fluorescence images and quantitative analysis of GFP in HEK293 cells after light-triggered CRISPR/Cas9 release from the liposomes are shown in Fig.2. When cells were treated with the liposomes alone, a slightly lower GFP fluorescence intensity was observed, compared with the control group without any treatment (about 5% less than the control), indicating the stability of the liposome formulation during incubation with the cells. With light illumination, CRISPR/Cas9 complex was released from the liposomes and knocked out the GFP, resulting in the clear reduction of its fluorescence signal. The lowest GFP expression level was achieved after 6 min illumination, compared with the liposome transfected cells without light irradiation (52.8% v.s. 94.8%). We also tested GFP gene knockout efficacy by employing Lipofectamine 2000 reagent as a delivery vehicle, for comparison purpose. The reduced GFP fluorescence intensity was observed in HEK293 cells at 48 hours after treatment. Although the similar GFP transfection effect was observed by using commercial lipofectamine, the on-demand gene release was achieved by using our light-triggered liposomes. This indicates that release of CRISPR system in a temporally controllable way would be possible by combining a delivery vehicle with light.

**Figure 2.**
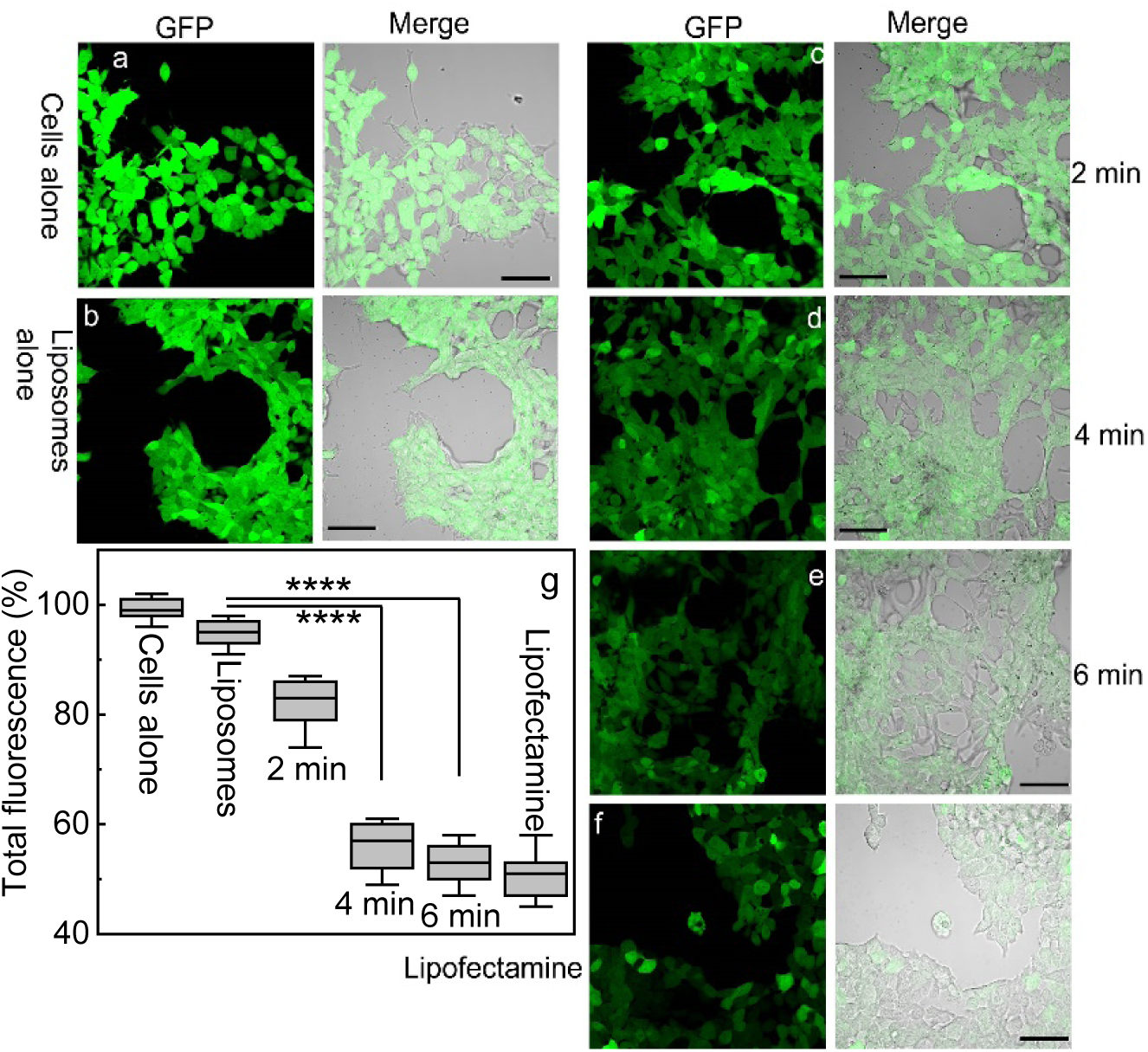
Confocal images of GFP expression in HEK293 cells without any treatment, (b-e) GFP expression after 48 hr of liposome transfection with and without light illumination and (f) GFP expression after 48 hr of lipofectamine transfection. Scale bars = 30 µm. (g) Quantitively analysis of GFP knockout efficiencies of different treatment groups. The box is bounded by the first and third quartile with a horizontal line at the median and whiskers extend to 1.5 times the interquartile range. The mean value was analysed using the t test (n=4). ****p <0.001, compared to the liposome group without light.

### A visual reporter system for rapid quantification of knockout efficiency *in vivo*

To assess the efficiency of light-sensitive liposome delivery of CRISPR/Cas9 *in vivo*, we developed a quantitative visual reporter system in zebrafish. Previously, we have shown a quantitative readout system *in vivo* to assess HDR-stimulating genome editing ^46^. Here, we established a gene knockout strategy where eGFP expressed specifically in the slow-muscle fibers of a stable transgenic zebrafish is knocked-out by CRISPR/Cas9 (Fig 3A). Slow-twitch muscle fibers form a single superficial layer directly under the skin arranged in parallel with the long axis of zebrafish (Fig 3B). Here we used a transgenic zebrafish line (smyhc1:eGFP) where slow-muscle specific smyhc1 promoter drives the eGFP expression at slow-twitch muscle fibers (Fig 3C). To generate a highly efficient DSB, we screened eight sgRNAs targeting eGFP using the reporter system and selected the sgRNA exhibiting highest rate (88.24%) of cutting efficiency (Sup Table 1). To assess the efficiency of the visual reporter system, we co-injected sgRNA targeting eGFP locus with Cas9 protein into single-cell zebrafish embryos (Fig 3D). We observed the loss of green fluorescent signal across individual slow-twitch muscle fibers showing loss of eGFP expression, whereas the control group injected without eGFP sgRNA did not exhibit any loss of green fluorescent signal (Sup Video 1 A-B). This allows rapid visual quantitation of knock-out efficiency at singe cell resolution *in vivo*.

**Figure 3.**
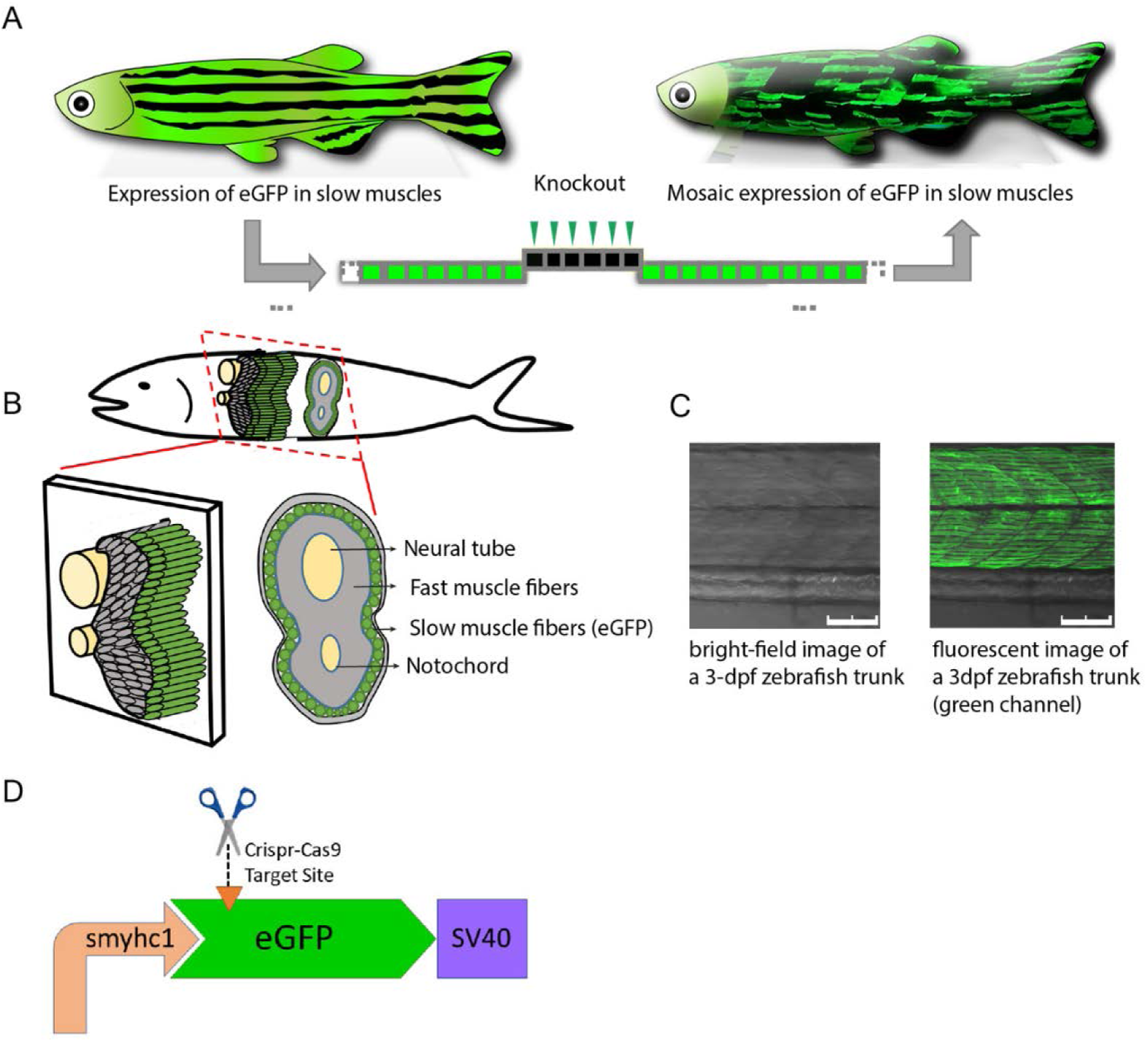
Schematic illustration of quantitative readout detection system in vivo. (A) Overview of the visual knock-out readout in zebrafish. (B) Schematic representation of zebrafish cross-section showing slow muscles forming a single layer of parallel fibers underneath the zebrafish skin. (C) Confocal section of smyhc1:eGFP zebrafish line under brightfield and green channel. Scale bars: 75 µm. (D) sgRNA-Cas9 complex targeting the eGFP expression driven by slow muscle-specific smyhc1 promoter.

### Assessment on *in vivo* knockout of eGFP gene by light-triggered liposomes

After confirmation of *in vitro* CRISPR transfection, we tested whether we can demonstrate targeted knockout of the eGFP by controlled release of CRISPR/Cas9 in zebrafish using light-triggered liposomes. To determine the effect of the light-triggered genome editing, transgenic smyhc1: eGFP zebrafish embryos were co-injected with Cas9 protein and liposomes encapsulating verteporfin and eGFP sgRNA. The injected embryos were randomly divided into two groups; either no light exposure or light exposure at 690 nm for 5 minutes. We used the visual reporter system described above to evaluate the efficiency of light-controlled genome editing *in vivo*. The initial qualitative assessment showed a major loss of green fluorescence signal in muscle fibers, suggesting light triggered release of CRISPR/Cas9 (Figure 4A, B). The negative control group did not show any loss of green fluorescent signal, highlighting the specificity of the assay. Therefore, we proceeded to quantify the total number of slow muscle fibers knocked out in the trunk of each embryo (n=80 embryos per group; Figure 4C, S4A). No light exposure resulted with modest but significant loss of green fibers compared to the negative control (Figure 4C; -ve control, 0 ± 0; light (-), 53.15 ± 35.38, p<0.0001, one-way ANOVA with multiple comparison). In contrast, embryos exposed to light activation showed a dramatically significant decrease in number of green fibers compared to no light control group, implicating light-triggered knockout of eGFP *in vivo* (Figure 4C; light (-), 53.15 ± 35.38; light (+), 308.37 ± 40.21, p<0.0001). Compared to positive control group injected with eGFP and Cas9 without liposome, embryos exposed to light activation showed a similar level of decrease in number of green slow-muscle fibers. We further observed that results from our quantitative model are consistent with the total fluorescence intensity results (Figure 4B, C).

**Figure 4.**
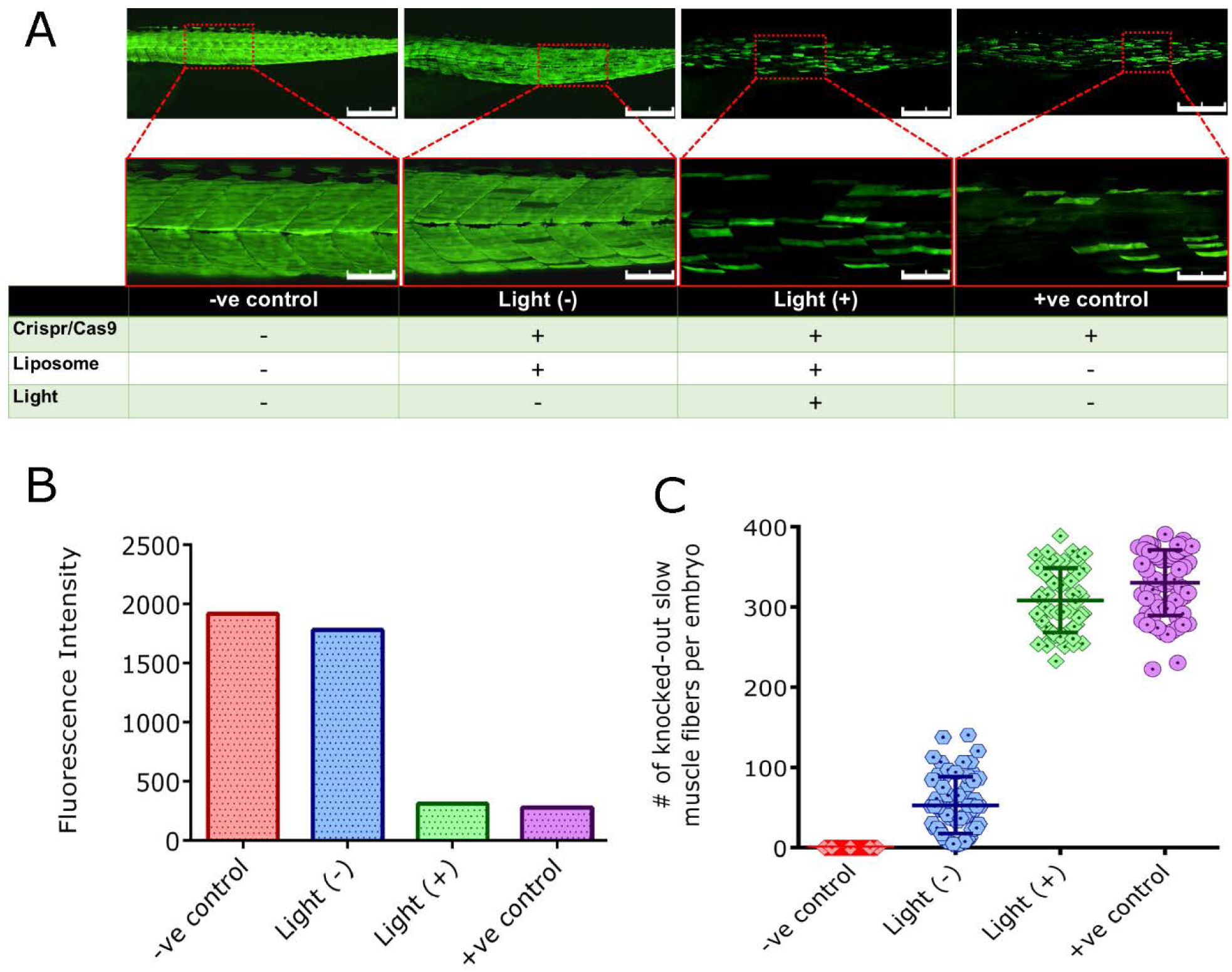
Light-triggered release of CRISPR/Cas9 in zebrafish. (A) Fluorescence images of smyhc1-eGFP zebrafish (3dpf); uninjected negative controls, co-injected with Cas9 and liposome/CRISPR complex without light exposure, co-injected with Cas9 and liposome/CRISPR complex with 5 min light exposure, and injected with only CRISPR/Cas9 as positive control and (B) Qualitative assessment of the knockout rate in zebrafish images by total fluorescence intensity. (C) Quantification of CRISPR/Cas9-mediated knockout rates in zebrafish by number of knocked-out slow-muscle fibers at single cell resolution. Scale bars: 500 µm, main image and 100 µm, inset.

To optimize the light-triggered release of CRISPR/Cas9 in zebrafish, we first compared different light exposure times using visual reporter system as a testbed. Embryos co-injected with liposome nanoparticles and Cas9 protein were subjected to one of five different irradiation times, 1 min, 2 min, 5 min, 15 min, 60 min. Qualitative assessment implicated a difference between light exposure times, suggesting longer exposure to light leads to higher knockout rates (Figure 5A, B). The quantitative analysis of the single-fiber analysis showed longer light exposure times leading to higher loss of green slow-muscle fibers. (Figure 5C; No Light, 27.84 ± 9.81; Light (1 min), 113.12 ± 14.77, Light (2 min), 300.67 ± 30.16 Light (5 min), 326.12 ± 36.55; n=60 embryos per group). However, we did not observe any significant difference in loss of green fluorescent signal at light illumination longer than 5 minutes (Figure S4B; Light (5mins), 326.12 ± 36.55; Light (60mins), 332.65 ± 33.60, p=0.43). Light illumination up to 5 min did not affect the embryo survival, however longer exposure to red light led to reduced embryo viability. At 60min light illumination, 36% of zebrafish morphologically normal looking zebrafish embryos remained alive (Sup Fig S5A).

**Figure 5.**
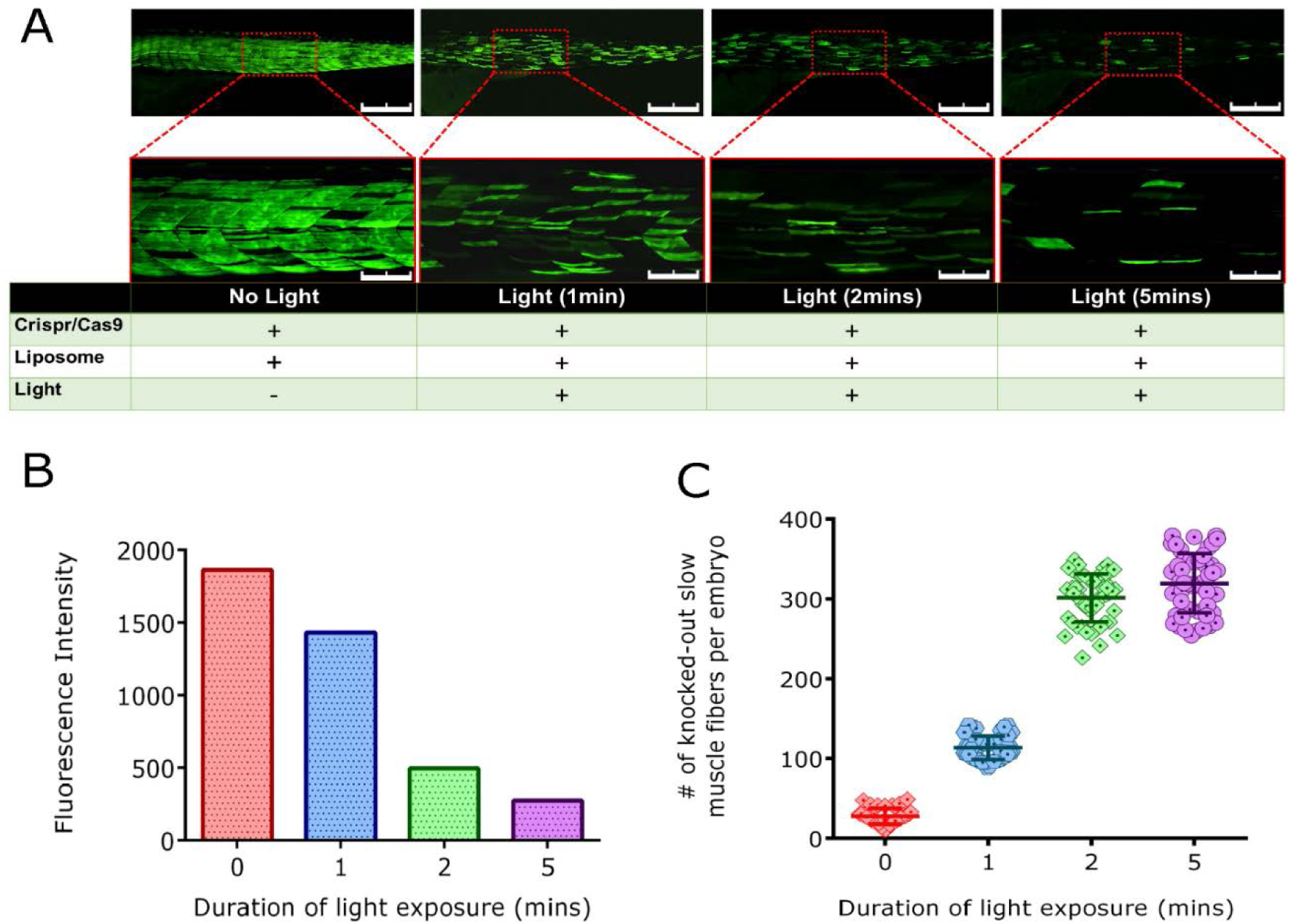
Effect of light exposure time on controlled release of CRISPR/Cas9. (A) Fluorescence images of smyhc1-eGFP zebrafish (3dpf) co-injected with Cas9 and liposome/CRISPR complex with no light exposure; 1 min of light exposure; 2 min of light exposure and 5 min of light exposure. (B) Qualitative and (C) quantitative assessment of the effects of light exposure times on the efficiency of CRISPR/Cas9-mediated knockout in zebrafish embryos. Scale bars: 500 µm, main image and 100 µm, inset.

Next, we investigated the effect of liposome nanoparticles concentration on light-triggered release of CRISPR sgRNA. We also determined the effect of liposome concentration on embryo toxicity by measuring the hatching rate of zebrafish embryos injected with different liposome concentrations. While the higher concentration of liposome led to increased mortality in zebrafish embryos (Figure S5B), efficiency of light-triggered release of CRISPR remained unaffected (Figure S6B).

## DISCUSSION AND CONCLUSIONS

The ability to manipulate any genomic sequence by CRISPR gene editing has created diverse opportunities for biological research and medical applications. However, further advancement of gene editing requires the development of optimal delivery vehicles ^8, 9, 15, 48-50^. Non-viral delivery is particularly advantageous, as it avoids insertional errors and it allows tight control over the dose, duration, and specificity of delivery ^9, 11, 15^. The liposomal platform investigated here is able to simultaneously release controlled amounts of the Cas9 nuclease and matching amounts of gRNA in a way that is spatially and temporally controlled by an external light beam applying safe levels of 690 nm light (0.15 mW/cm^2^) to tissue surface. While light required to trigger our liposomes penetrates tissue only up to a few millimeters ^51^, optical fibre approaches developed for photodynamic therapy of cancer make it possible for these liposomes to be applied in deep tissue as well ^52^.

Light-triggered liposomal release of CRISPR reagents offers previously unavailable option for gene editing to be localised in space and time; such four-dimensional control will be important for novel research applications and for further clinical translation of the CRISR-Cas9 technique. Lipid nanoparticles and conventional liposome-based delivery widely used for CRISPR transfection in preclinical settings suffer from a drawback. After internalization of the through the endocytic pathway, most of these carriers become entrapped in endo/lysosomes where the enzymatic degradation may result in deactivation of CRISPR components before they are able to be released to perform their gene editing action ^53^. Therefore ensuring rapid endo/lysosomal escape of the cargos is required for efficient CIRSPR/Cas9 transfection via lipid-based nanoparticles ^54^. Our light-triggerable liposomes overcome the issue of endo/lysosomal entrapment, because, as we established earlier, VP activated by light illumination generates sufficient singlet oxygen to destabilise not only the liposomes but also the liposomal and endo/lysosomal membranes ^30^. We demonstrated this by encapsulating antisense oligodeoxynucleotides (asODN) in this platform and quantitative assessment of the endo/lysosomal escape based on the released profiles of DNA molecules and endo/lysosomes ^30^. This delivery system was shown to enable an effective knockdown of target gene (PAC1R, 74 ± 5% reduction) and inhibition of the neurite-growth of PC12 cells via PACAP-dependent signaling pathway.

The ability of our liposomes to deliver defined amounts of intact Cas9 represents a key advantage of this formulation for efficient and nontoxic gene editing. The Cas9 protein is large (∼160 kDa) and this prevents its direct delivery to cells ^48^. We found in this work that our liposome encapsulation enables direct Cas9 protein delivery to cells and may partially protect it from degradation. Such direct nuclease delivery in CRISPR offers the immediate function without protein expression process and the most rapid therapeutic activity as there is no cellular translation or transcription ^4^. Direct delivery of purified nuclease proteins or Cas9 protein-gRNA complexes is additionally important because it yields high levels of gene editing ^55^. This is consistent with the results reported here of high efficiency of the *eGFP* knockout observed in HEK293 cells (up to 52%) and in zebrafish embryo (up to 77%) treated with light-triggered liposomes compared to the control group (Fig. 2 and 4). Our result confirms that light-triggered CRISPR/Cas9 release does not compromise the genome editing activity in the target loci. Transient protein delivery via liposomes also restricts the duration of nuclease activity potentially reducing off-target editing as the nuclease has less opportunity for promiscuous action ^10^. Our approach may therefore play an important role in ensuring precision and safety of the CRISPR-Cas9 tools. The liposomes also enable direct gRNA delivery to cells which is not straightforward because the long phosphate backbone of gRNA is too negatively charged to passively cross the membrane. Furthermore, the liposomes may help the gRNA to avoid nuclease degradation. We found in this work that our liposome encapsulation provide sufficient protection for CRISPR reagents to gain cellular entry in HEK293 cells and in zebrafish embryos and subsequently escape from the endosomes to enter the cytoplasm while remaining functional ^9, 48^. We also observed only a modest leakage of liposomal contents in controls which were not exposed to light. This is tentatively explained that the cell contents may compromise the integrity of liposomal membranes ^56, 57^. The liposomal nanoparticles demonstrated minimal cytotoxicity both in HEK293 cells and zebrafish embryos, under the current experimental conditions (Fig. S2).

The data shown in Fig. 2 compare our light-triggered liposomal delivery with CRISPR delivered using Lipofectamine, a commercially available liposome delivery vehicle for nucleic acids and gene editing proteins. Lipofectamine draws on the ability of lipids to spontaneously form nanoparticles in aqueous solution in order to protect their hydrophobic tails from the solvent. By simple mixing, a payload may be encapsulated within a lipid nanoparticle. Lipofectamine contains cationic lipids that complex with the negatively charged nucleic acid molecules and this reduces the effect of electrostatic repulsion of the negatively charged cell membrane ^58, 59^. This additionally protects nucleic acids from nucleases and allows them to be taken up by target cells. Lipofectamine has been previously used in conjunction with the CRISPR system for various application purposes, including generation of an immunodeficiency model ^60^, multiplex genome editing ^61^, and gene therapy of cystic fibrosis and bladder cancer ^62, 63^. The *in vitro* CRISPR transfection efficiency via our light-triggered liposomes and Lipofectamine was found to be comparable (52% v.s. 50% GFP level reduction in Fig. 2). However, unlike Lipofectamine, our liposomes can be triggered by light allowing spatial and temporal control of gene editing, moreover they are feasible to be functionalized with different ligands of interests

The light-triggerable CRISPR delivery vehicles reported here are biocompatible and made entirely from clinically-approved components using a simple synthesis method. This design avoids the need for numerous manufacturing steps in the future scaling-up process. It is also important from a commercial and regulatory point of view that the entire gene therapy product can be packaged in a single vehicle. *In vivo* gene editing benefits from tissue-specific targeting (e.g. using tissue specific promoters of Cas9) to prevent undesirable off-target gene editing events. Targeted delivery of liposomes is well established ^26^, and such molecular targeting is also directly applicable to the CRISPR-carrying liposomes investigated here. Liposomes are also well suited to co-delivery of multiple components, and this is highly relevant as novel CRISPR refinements may require simultaneous delivery of multiple functional entities. The liposomes are entirely DNA-free and this will help avoid DNA toxicity and stimulating immune responses. Favourable biodistribution in specific disease conditions may be achieved by optimising formulations and by a suitable route of administration. Spatial and temporal control of gene editing using the liposomal delivery vehicles reported here will open new options for exciting science and wider translation of CRISPR-Cas9 gene editing.

## ASSOCIATED CONTENT

### Supporting information

The Supporting Information is available free of charge:

- Experimental setup of the cell work, cell’s viability assessment, 3D rendered confocal images of individual slow-muscle fibers expressing eGFP and quantitative assessment in zebrafish by counting the number of knock-out slow muscle fibers per embryo (PDF).
- Fluorescent signal of eGFP across individual slow-twitch muscle fibers (Video)

## AUTHOR INFORMATION

### Notes

The authors declare no competing financial interest.

## ACKNOWLEDGEMENTS

We acknowledge Prof. Paul Pilowsky from Heart Research Institute, Sydney, Australia for drawing our attention to the problem of spatial and temporal control of gene engineering. of gene editing research. All TEM images in this work were performed in the Microscopy Unit, Faculty of Science and Engineering at Macquarie University. This work is partially supported by the ARC awards CE140100003 and DP170101863.

## AUTHORS’ CONTRIBUTIONS

W. D. and Y. A. conducted the experiments and drafted the manuscript. W. C. synthesized the liposomes and provided the TEM image. E. G. designed and supervised studies as well as reviewed and revised the manuscript.

