## Supporting figures and tables for "Spatial and temporal control of CRISPR/Cas9-mediated gene editing delivered via a light-triggered liposome system"

Supporting information

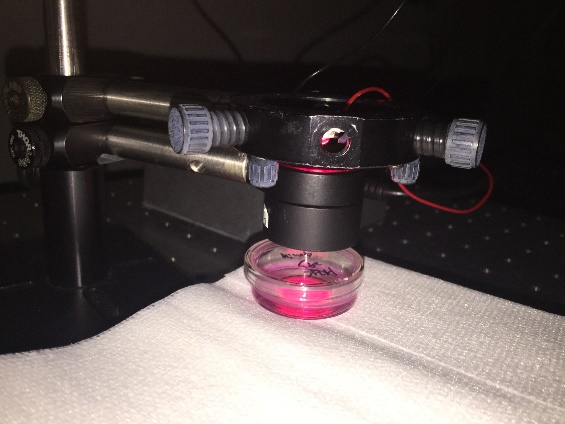

**Figure S1** Experimental setup of cells under a LED illumination at 690 nm.

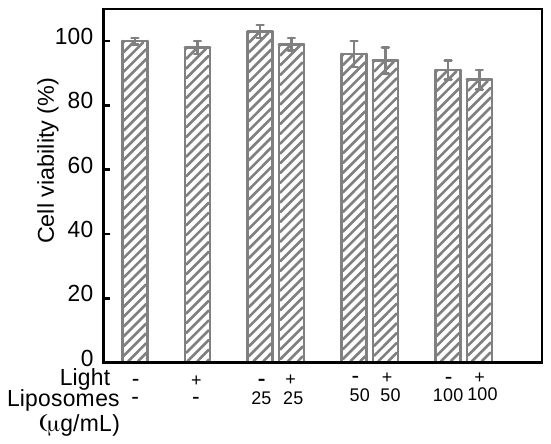

**Figure S2** The viability of HEK293 cells after incubation with the liposomes for 4 hr and illumination at 690 nm for 6 min. The concentration of liposomes was 25, 50 and 100 µg/mL. The viabilities are expressed as mean percentages and standard deviation (n=4) relative to control cells.

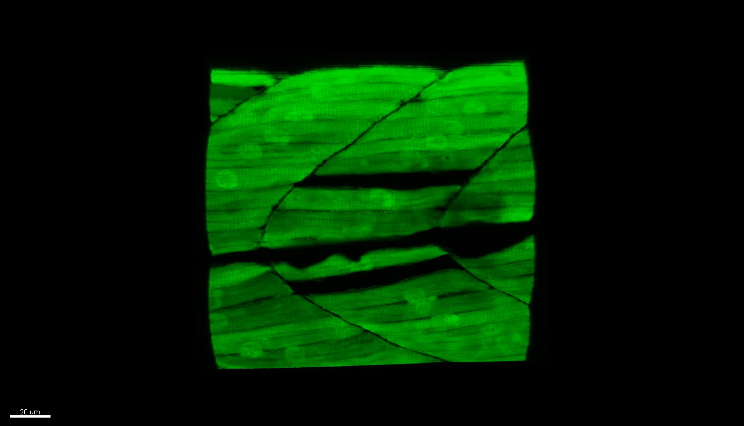

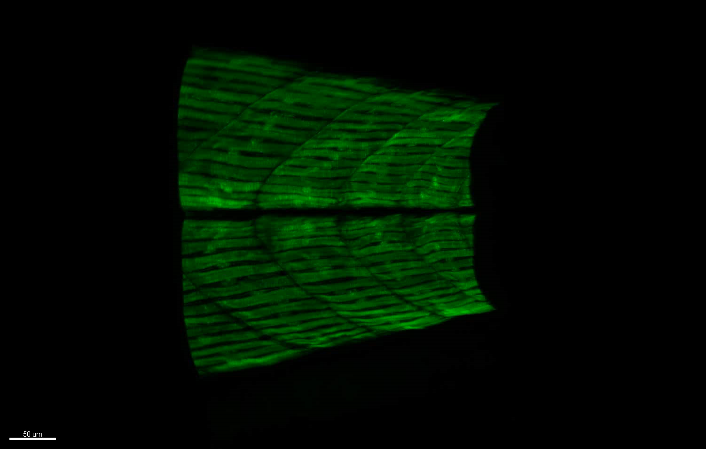

B

A

**Figure S3** 3D rendered confocal images of (A) control transgenic smyhc1:eGFP embryos show individual slow-muscle fibers expressing eGFP as a single layer and (B) transgenic embryos injected with eGFP-targeting CRISPR guide RNA show loss of green fluorescent signal from slow-muscle fibers

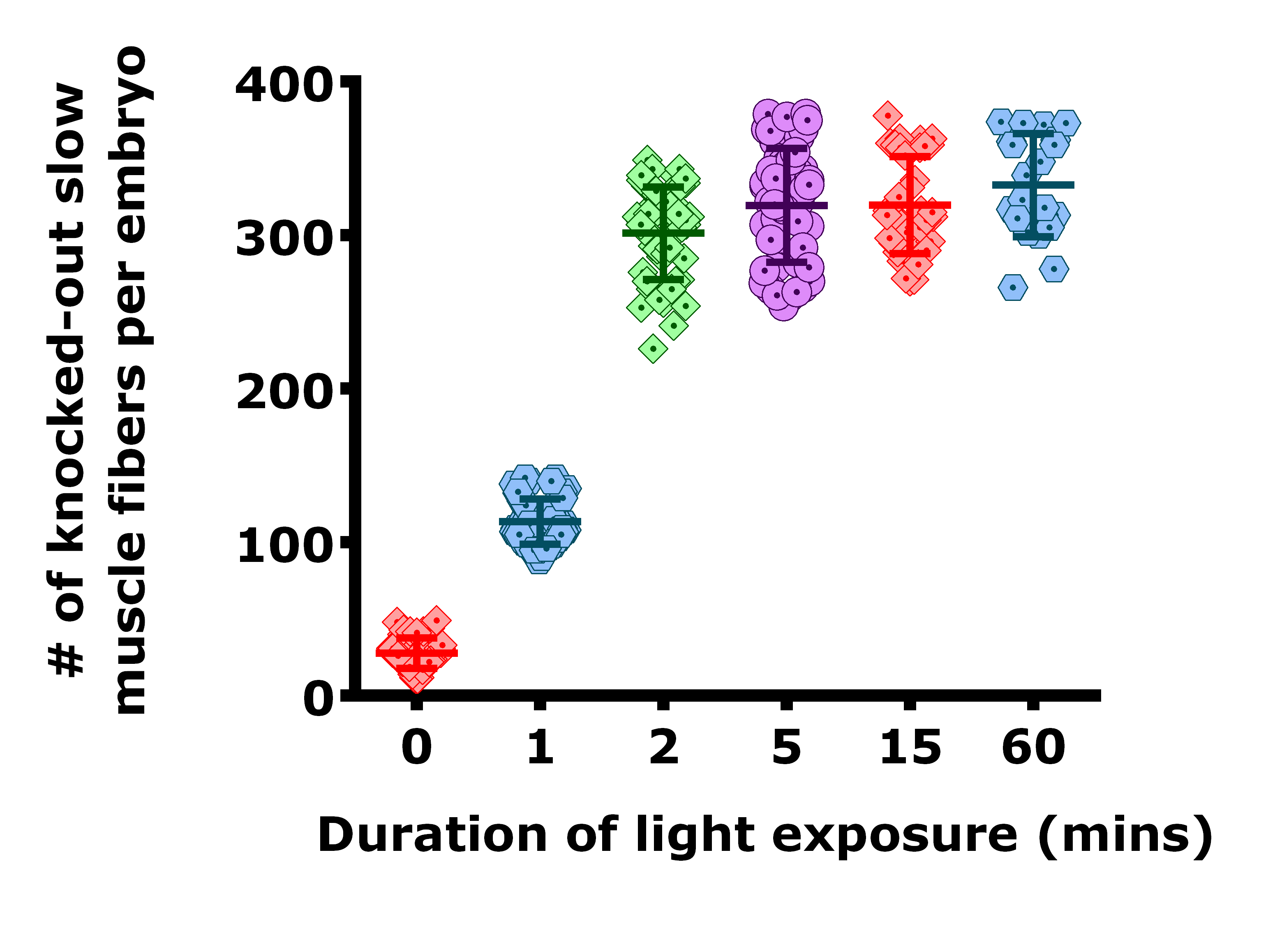

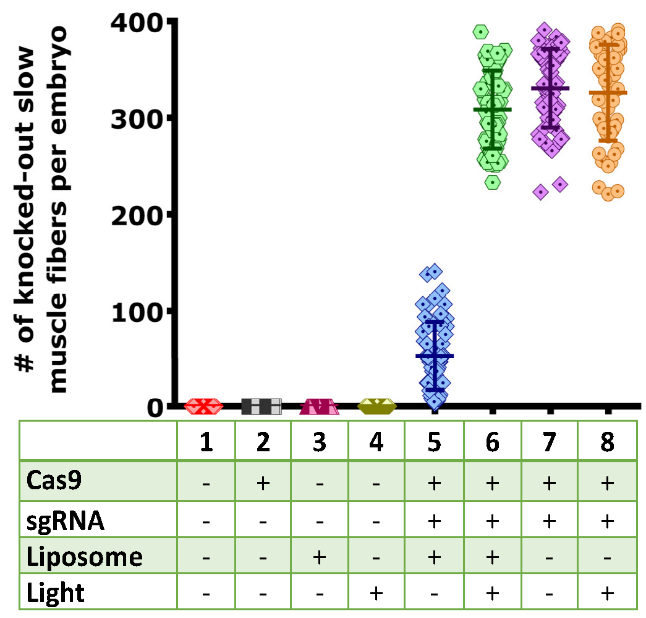

A

B

**Figure S4** (A) Quantitative assessment of Cas9-mediated knockout by light-triggered release of CRISPR in zebrafish by counting the number of knock-out slow muscle fibers per embryo under different treatment as indicated in the image; (B) Effect of light exposure time on controlled release of CRISPR/Cas9 by counting the number of knock-out slow muscle fibers per embryo at the different illumination time points

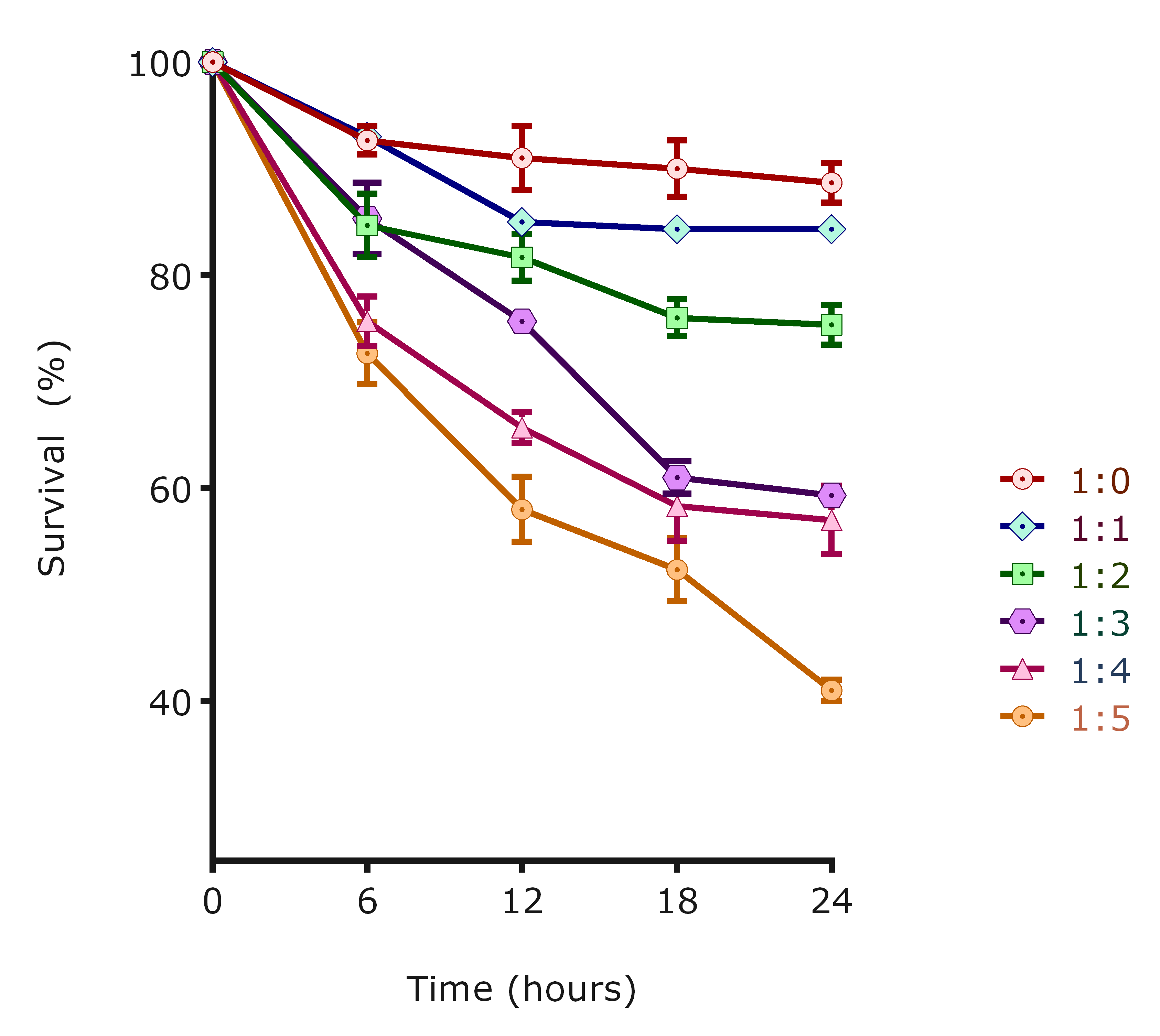

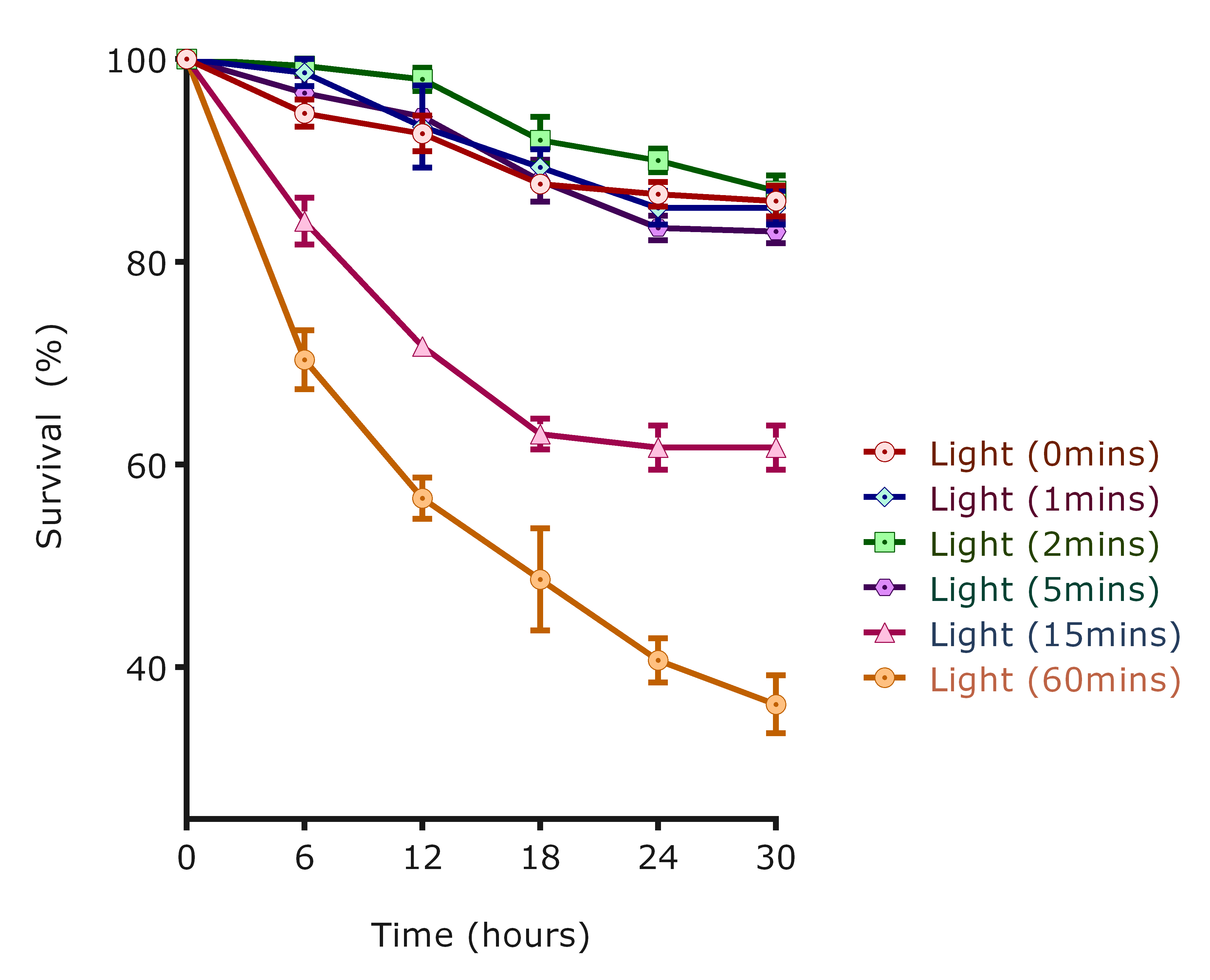

B

A

**Figure S5** Assessment of light and liposome toxicity to zebrafish embryos. (A) Survival of 3dpf zebrafish injected with different CRISPR/Cas9 to liposome concentrations. (B) Survival rate of zebrafish embryos exposed to different duration of light.

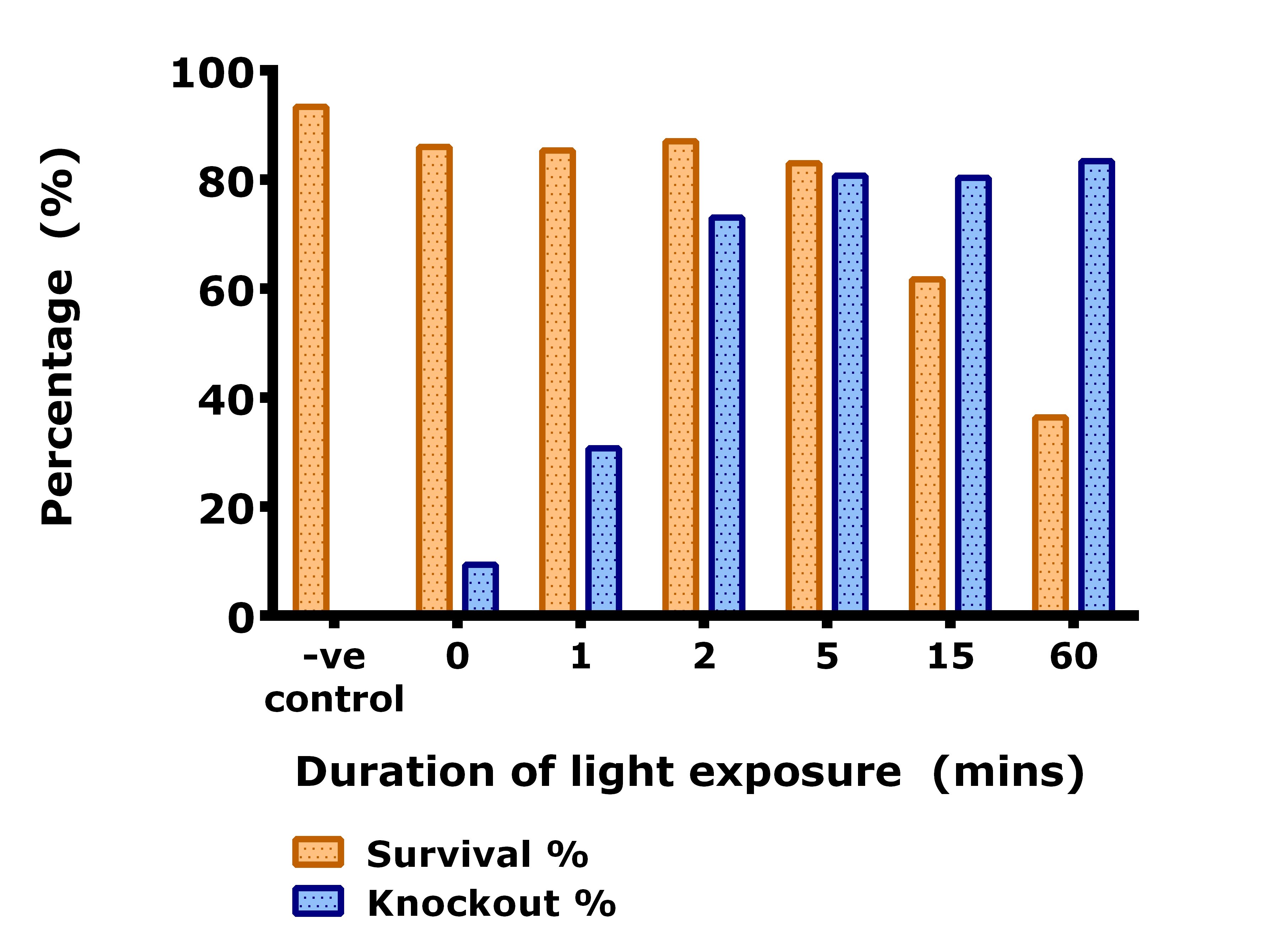

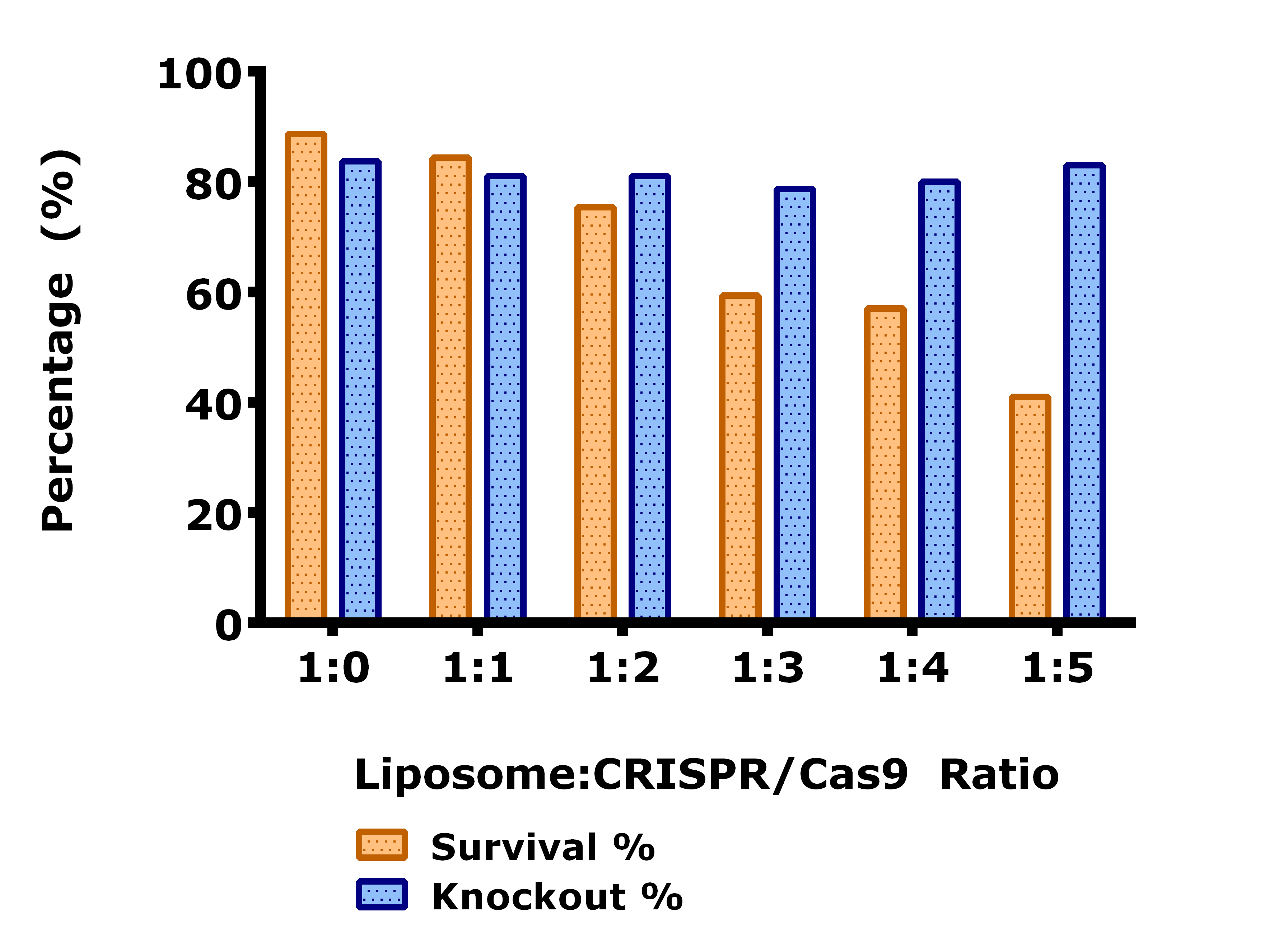

**Figure S6** The survival and knockout rates (%) of zebrafish embryos (A) exposed to different duration of time (between 0 - 60 mins) and (B) injected with different CRISPR/Cas9 to liposome ratios (between 1:0 - 1:5)

| **ID** | **Sequence** | **Target** | **Cutting Efficiency (%)** |
| --- | --- | --- | --- |
| sg_eGFP_01 | ATGGTGAGCAAGGGCGAGG | eGFP | 37.80 |
| sg_eGFP_02 | GACCAGGATGGGCACCACCCCGG | eGFP | 56.18 |
| sg_eGFP_03 | CGCCGGACACGCTGAACTTGTGG | eGFP | 6.86 |
| sg_eGFP_04 | CAAGTTCAGCGTGTCCGGCGAGG | eGFP | 26.33 |
| sg_eGFP_05 | GGCGAGGGCGATGCCACCTACGG | eGFP | 66.04 |
| sg_eGFP_06 | GGGCACGGGCAGCTTGCCGGTGG | eGFP | 88.24 |
| sg_eGFP_07 | AGCACTGCACGCCGTAGGTCAGG | eGFP | 50.80 |
| sg_eGFP_08 | GCTTCATGTGGTCGGGGTAGCGG | eGFP | 49.10 |

**Table S1** Comparison of cutting efficiency of CRISPR gRNA targeting eGFP gene. Cutting efficiency of eight sgRNAs targeting different loci on eGFP gene was determined using RFLP and further validated by observing the loss of green fluorescence signal from slow-muscle fibers.
